# Masked superstrings as a unified framework for textual *k*-mer set representations

**DOI:** 10.1101/2023.02.01.526717

**Authors:** Ondřej Sladký, Pavel Veselý, Karel Břinda

## Abstract

The popularity of *k*-mer-based methods has recently led to the development of compact *k*-mer-set representations, such as simplitigs/Spectrum-Preserving String Sets (SPSS), matchtigs, and eulertigs. These aim to represent *k*-mer sets via strings that contain individual *k*-mers as substrings more efficiently than the traditional unitigs. Here, we demonstrate that all such representations can be viewed as superstrings of input *k*-mers, and as such can be generalized into a unified framework that we call the masked superstring of *k*-mers. We study the complexity of masked superstring computation and prove NP-hardness for both *k*-mer superstrings and their masks. We then design local and global greedy heuristics for efficient computation of masked superstrings, implement them in a program called KmerCamel, and evaluate their performance using selected genomes and pan-genomes. Overall, masked superstrings unify the theory and practice of textual *k*-mer set representations and provide a useful framework for optimizing representations for specific bioinformatics applications.

## 1 Introduction

The emergence of modern DNA sequencing methods resulted in exponentially growing amounts of sequence data that are becoming increasingly difficult to analyze using the existing computational techniques [1]. One approach to address the ever-increasing abundance of sequences has been a shift towards *k*-mer-based methods that allow for substantially more efficient analyses in big data genomics compared to the traditional methods based on alignments. Examples of problems where *k*-mer-based methods play an important role include large-scale data search [2, 3, 4], metagenomic classification [5, 6], RNA-seq abundance estimation [7, 8], or infectious disease diagnostics [9, 10] to mention at least a few.

However, the shift towards *k*-mers introduced new computational challenges, such as how to represent large *k*-mer sets in random access memory (RAM) and on disk, and in connection to the individual *k*-mer-based data structures [11]. While *k*-mers can easily be encoded as numerical values, which is widely used for instance in *k*-mer counting [12, 13], such representations are unable exploit the overlapping structure of the *k*-mers present in typical datasets that is usually referred to as the so-called spectrum-like property (SLP) of *k*-mer sets [14]. More compact approaches of *k*-mer set representations were developed later, incorporating SLP as an implicit structural assumption and using it to compact *k*-mers overlapping by *k* − 1 nucleotides (review in [15]). First of them were unitigs [16, 17], which can be seen as non-ambiguous contiguous sequences in the data*’*s de Bruijn graphs; unitigs had been previously well-studied and implemented in other contexts, such as genome assembly, and were thus easy to use.

However, unitigs tend to impose large computational overheads due to the requirement of stopping at all branching nodes in the underlying de Bruijn graphs [18, 19]. This is particularly pronounced under in the presence of extensive biological variation, such as in bacterial pan-genomics. Further improvements focused on increasing the efficiency and scalability of *k*-mer set representations by relaxing the stopping constraint – first by simplitigs/Spectrum Preserving String Sets [20, 18, 19] (and eulertigs [21] as the optimal form of this representation) and later also by matchtigs [22], which further relaxed the requirement of *k*-mers having to occur exactly once in the representation that was blocking additional compaction.

However, all of these representations are limited by their reliance on the availability of (*k* − 1)-long overlaps between the *k*-mers to be represented. In fact, many situations such long overlaps do not exist or cannot be fully exploited. One example is provided by *k*-mer sets arising from subsampling – in such a case, even matchtigs, which is the most relaxed existing textual representation, would necessary contain a huge number of very short strings, including many individual isolated *k*-mers, which is an undesirable property. Moreover, modeling of *k*-mer set representations via sets of strings whose length is to be minimized, does not fully align with the computational reality, in which the management of individual sequences always incurs substantial overheads (see, e.g., [18, Fig 6b, memory]). For instance, in the common implementations of text indexes, such as the FM-index [23] in BWA [24], all strings are usually concatenated together and the index then maintains an auxiliary data structure to keep track of their starting positions in the concatenation, leading to a substantial performance degradation when the sequences are too numerous.

Here we propose the concept of *masked superstrings*, combining the idea of representing *k*-mer sets via a string that contains the *k*-mers as substrings, with masking out positions of the newly emerged *“*false positive*” k*-mers. This allows to unify the notions of existing representations, remove their main limitation of using (*k* − 1)-long overlap only, and provide novel ways how to represent *k*-mer data more compactly. We thoroughly evaluate the whole concept of masked superstrings, including the associated computational complexities, implementation strategies, and performance on real data using a newly developed tool called KmerCamel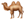. Throughout the paper, we demonstrate that masked superstrings provide a useful building block for future *k*-mer-based data structures.

## 2 Setup and preliminaries

We consider the standard setup of the problem of *k*-mer set textual representations [15]. We assume that *k* is a positive integer and *K* is a set of *k*-mers to be represented. We consider the bi-directional version of the setup, in which *k*-mer and its reverse complement are considered a single mathematical object, with the exception of illustrations and examples that are always provided in the uni-directional model for simplicity.

To unify the terminology throughout the paper, we call the simplitig/SPSS representation [20, 18, 19], which does not allow multiple occurrences of the same *k*-mer, simply *SPSS* (*Spectrum-Preserving String Set*) and call its members *simplitigs*. For the more general matchtig representation [22], which does allow multiple occurrences of the same *k*-mer, we use the term *rSPSS* (*Repetitive Spectrum-Preserving String Set*) and for its members the term *matchtigs*. For a reference to either of these representations, we use simply the term *(r)SPSS*.

Fig. 1a,b depicts an example of a *k*-mer set *K* and the corresponding de Bruijn and overlap graphs (see Appendix B for definitions). Fig. 1c shows the corresponding examples of the state-of-the-art (r)SPSS representations of *K* (rows 1-4 in the *‘*Strings*’* column).

**Figure 1:**
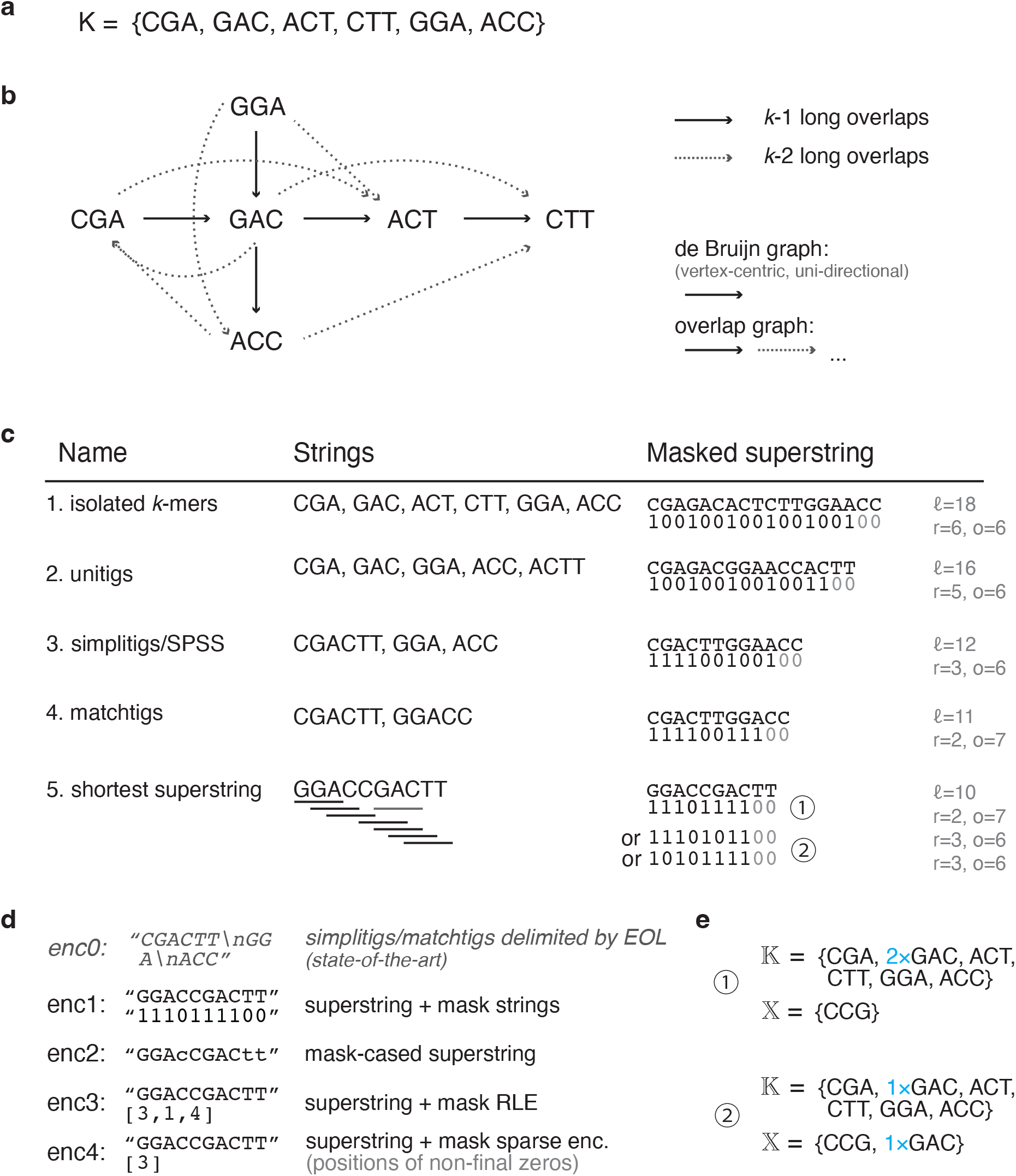
Overview of the masked superstring concept for representing *k*-mer sets. We focus on the uni-directional model for simplicity. **a)** An example of a *k*-mer set (*k* = 3). **b)** The corresponding de Bruijn (solid edges) and overlap graphs (solid and dashed edges, respectively). **c)** Individual representations of the *k*-mer set and the corresponding masked superstring, sorted with respect to the length. Value *ℓ* is the superstring length (generalizing the cumulative length in (r)SPSS), *r* is the number of runs of ones (generalizing the number of sequences in (r)SPSS), and *o* is the total number of ones in the mask. **d)** Examples of the encodings the shortest superstring. **e)** The represented and ghost *k*-mer multisets of the superstring from c5 (𝕂 and 𝕏, respectively). Note that the individual strings in c1–c4 could be concatenated in different orders, which would result in different sets of ghost *k*-mers, as well as change the multiplicities in 𝕂.

## 3 Results

### 3.1 The concept of masked superstrings

We developed *masked superstrings* as a generalizing concept, comprising the state-of-the-art textual representations for *k*-mer sets that follow the spectrum-like property (SLP) [14] (a structural assumption characterizing typical *k*-mer sets encountered across bioinformatics in mainstream applications), allowing better compression capabilities, and also adding support for other types of *k*-mer data that do not follow SLP.

We built upon the observation that the state-of-the-art (r)SPSS representations are deeply rooted in to the concept of superstring [18, 15], which has been deeply studied in many areas of computer science (for a primer of the shortest superstring problem, see Appendix B).

The core idea behind masked superstrings is shifting from a set of strings, whose *k*-mers are *exactly* the *k*-mers to be represented, to *some* superstring of the *k*-mers, with the goal of their better compaction that can be achieved through the relaxation of restrictions on the lengths of *k*-mer overlaps.

However, as such a superstring contains also other, *“*false positive*” k*-mers (we will call them ghost *k*-mers) that act as noise, the superstring needs to be further equipped with a binary mask to signalize which *k*-mers are in fact to be used and which are to be ignored. We summarize the relation between *K* and masked superstrings (*M, S*) in the following definition.

#### Definition 3.1.

*Given a positive k-mer length *K* and a k-mer set K, a* masked superstring *of K is any pair of strings S and M of the same length ℓ, over the ACGT and binary alphabets, respectively, satisfying*

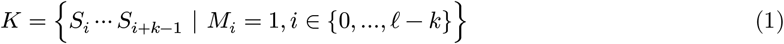

*We call S and M the k*-mer superstring *and its* mask, *respectively. By convention, we set the last, unused k* − 1 *mask bits to zero*.

We make two remarks about the practical aspects of masked superstrings. First, the *k*-th mask position from the right will always set to one in practical applications, therefore, such as a mask encodes also the *k*-mer length *k* (by counting the *k* − 1 zeros from the right). Second, while we define the masked superstring representation using two strings, *𝒮* and *M*, for practical purposes, it is often convenient to combine them into a single string, called *mask-cased superstring*, over the ACGTacgt alphabet, where uppercase letters represent 1 in the mask and lowercase letters correspond to 0 (see enc2 in Fig. 1d).

### 3.2 Relation of masked superstrings to (r)SPSS and their associated algorithms

Masked superstrings naturally generalize the concept of (r)SPSS. To see that, we note that the problem of representing *k*-mer sets always involves two major aspects: *“*How can *k*-mer sets be represented as compact mathematical objects?*”* and *“*How can these representations be computed efficiently?*”* In the case of (r)SPSS, the compact mathematical object is a set of strings whose *k*-long substrings form exactly *K*, and the efficient algorithms are greedy methods such as ProphAsm [18], UST [19], or greedy matchtigs [22].

Both of these (r)SPSS concepts can be directly translated to the realm of superstrings through the following relations.

**Algorithms: Every (r)SPSS algorithm is an approximation of the shortest** *k***-mer superstring**. Given a *k*-mer set *K* and an associated ordered (r)SPSS, the concatenation of its strings is a superstring of *K*. Moreover, when omitting artificial edge-cases that would never be encountered in practice, the total length of the superstring is no bigger than the number of *k*-mers (|*K*|) multiplied by *k* (a superstring with such length corresponds to concatenating all *k*-mers together, which is the most naïve and wasteful approach). As the resulting superstring length is always bounded from below by |*K*|, every such superstring of an (r)SPSS is necessarily a *k*-approximation of the optimal solution.

**Representations: Every (r)SPSS defines a masked superstring (but not conversely)**. Transforming an (ordered) (r)SPSS to a masked superstring is straightforward. Indeed, we just need to accompany individual strings by their corresponding masks and then concatenate the strings and masks in the same order. Specifically, for every string in an (r)SPSS, its corresponding mask is set to all 1s, with the exception of the last *k* − 1 characters that are set to 0s.

On the other hand, a general masked superstring does not translate to an (r)SPSS in the form of a split of the superstring (unless all runs of 0s in the mask are of the length *k* − 1). We illustrate the translation of (r)SPSS to masked superstrings in Fig. 1c and further characterize their relations in Tab. 3 in Appendix A.

### 3.3 Properties of superstring masks

While the only primary characteristic of a *k*-mer superstring to be optimized is its length *ℓ*, which is the direct generalization of the cumulative length (or weight) of simplitigs or matchtigs [18, 19, 22], *k*-mer masks have much more complex characteristics, and the choice of the one to optimize will depend on the specific application.

In particular, the same superstring *𝒮* of a *k*-mer set *K* can have many different masks *M*; the only requirement is that Equation 1 holds. In fact, for any *k*-mer that occurs multiple times, we can set to 1 any non-empty subset of its occurrence positions in the mask; therefore, the set of admissible masks forms a semi-lattice structure. Importantly, the specific choice of *k*-mer occurrences to include can impact both the frequencies in the multiset of represented *k*-mers 𝕂, as well as the ghost set *X* and multiset 𝕏 (Fig. 1e), which might have a substantial impact on the performance in downstream applications.

Furthermore, unlike superstrings that are expected to have a high entropy and thus being little compressible beyond 2 bits per *k*-mer, zeros in masks tend to be rather sparse in most *k*-mer sets, and various efficient encodings with little space requirements can be envisioned (see Fig. 1d for examples). Namely, we consider the following complexity measures related to these encodings:

- **Length of the mask** (*ℓ* in Fig. 1c), which models the space requirements of the mask stored as a bit-vector (enc1 in Fig. 1d).
- **Number of ones** (*o* in Fig. 1c), which models the space requirements of a sparse encoding of the mask (enc4 in Fig. 1d).
- **Number of runs of ones** (*r* in Fig. 1c), which models the space requirements of its run-length encoding (RLE) (enc3 in Fig. 1d).
- **Size after compression** using a general-purpose method such as xz (implementing the Lempel-Ziv Markov Chain Algorithm, available from http://tukaani.org/xz/) or RRR [25].

The *o* and *r* quantities generalize the number of sequences in the (r)SPSS representations with respect to different data structure implementations, and in addition to that, *r* can be seen as a first-order proxy to the mask compressibility.

### 3.4 Complexity of masked superstring computation

The computation of masked superstrings is a complex optimization problem that can be configured for individual bioinformatics applications by adjusting the objective function, so that it reflects the memory footprint of the associated indexes, computational complexity of operations of the associated algorithms, or other similar quantities. However, such a problem is difficult to tackle in its full generality.

Instead, we propose a simplified protocol, based on a two-step optimization procedure, starting by the computation of a superstring and followed by mask optimization; i.e., superstring and mask computation are considered two separate problems and thus can be studied independently of each other. Despite this simplification, we prove the NP-hardness for both of the problems.

### *k*-mer superstring computation is NP-hard

While the problem of shortest common superstring is known to be NP-hard in many different formulations (see Appendix B), to the best of our knowledge, this has never been formally shown for the problem of sets of canonical *k*-mers. Nevertheless, we found that NP-hardness holds even in this case and proved it by a reduction from a special case of the superstring problem; see Appendix F (Theorem F.1).

### Mask optimization can be anywhere between linear and NP-hard, based on the objective function

Unlike superstrings, masks can be optimized for a multitude of mutually conflicting measures, such as the minimum number of ones, maximum number of ones, or minimum number of runs. While for the former two cases, optimal masks are easy to compute in linear time (e.g., with the use of *k*-mer counting using 1 or 2 passes, respectively), the minimization of the number of runs is in fact NP-complete as we prove in Appendix G by a reduction from Set Cover (Theorem G.1), which actually shows a strong hardness of approximation for this mask optimization task.

### 3.5 Heuristics for superstring computation and mask optimization

To compute masked superstrings rapidly, we develop several greedy heuristics that generalize the concepts already studied in the literature and modify them for the bi-directional model.

#### Local vs. global graph exploration

We make a distinction between two fundamentally different approaches for superstring computation that appear in the literature under the roof category of greedy algorithms: *global greedy algorithms* and *local greedy algorithms*. While the former algorithms are based on merging individual strings while considering all strings at the same time (i.e., having a global view on the up-to-date overlap graph), the latter algorithms only operate with one string at a time that is being progressively extended locally (i.e., making a traversal through the overlap graph). The most famous example of a global greedy approach is the well-studied Greedy algorithm for approximately shortest superstrings (see e.g. [26, 27]), whereas ProphAsm [18] for simplitigs provides an example of a local approach. Here, we generalize both types of approaches for the use case of masked superstrings in the bi-directional model.

#### Global greedy

We developed a modified version of the traditional Greedy algorithm, tailored for *k*-mers in the bi-directional model that we call BiDir-GlobalGreedy (Appendix D). Canonical *k*-mers are accounted for by doubling *k*-mers in the underlying overlap graph and prohibiting edges between them.

#### Local greedy

We generalized the ProphAsm algorithm [18, Alg. 1] to smaller overlaps to obtain an algorithm that we call BiDir-LocalGreedy (Appendix C). In an iterative fashion, the algorithm selects an arbitrary *k*-mer as a seed of a new superstring segment and keeps extending it forwards and backwards at the same time as long as possible, by maximum overlap, while progressively increasing the extension step *d* in case no overlap of a given length was found, up to a predefined threshold *d*_*max*_. The already used *k*-mers are immediately removed from *K*. The extension proceeds by brutal force, first testing all four nucleotides, then testing all combinations of two of them, etc., while testing for the presence of the created *k*-mer. This process is repeated until all *k*-mers are covered. Note that setting *d*_*max*_ = 1 corresponds to the original ProphAsm.

#### Mask optimization using linear programming

Although the minimization of the number of runs is an NP-complete problem (Sec 3.4), real-life instances of similar problems tend to be far from the worst-case scenarios, and often can be *“*attacked*”* by a reduction to a well-studied optimization problem and a subsequent use of highly optimized solvers. Following this approach, we model the minimization of the number of runs as an integer linear program (ILP) and run an efficient ILP solver. The key idea is to find all maximal possible runs of ones in a mask for the given superstring and then select the minimum number of these runs such that all *k*-mers are covered, i.e., appear in at least one selected run. The resulting ILP is reminiscent to the Set Cover ILP. We also design a simple linear-time greedy heuristic for mask pre-optimization that reduces the size of the ILP significantly; see Appendix H for details.

### 3.6 Implementation

We implemented the two superstring heuristics in a program called KmerCamel 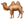, which first reads a user-provided FASTA file with genomic sequences, computes the corresponding *k*-mer set, computes a masked superstring using a user-specified heuristic and core data structure, and prints it in the enc2 encoding.

KmerCamel 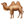 was developed in C++ and is available under the MIT license from Github (https://github.com/GordonHoklinder/kmercamel). Each of the two heuristics was implemented using two distinct data structures: a hash table and the Aho-Corasick automaton. The hash-table-based implementations follow the greedy algorithms as described in Appendices C (local) and D (global). The implementations of the two *k*-mer superstring heuristics using the Aho-Corasick automaton are described in Appendix E.

For the mask optimization, we implemented the greedy minimization and maximization of ones in custom Python scripts. For minimizing the number of runs, we created a transformation script that reads a provided masked superstring, builds the corresponding ILP problem according to Appendix H, and solves it using the PULP solver [28]. All the scripts are provided as a part of the evaluation pipelines in the Github supplementary repository (https://github.com/karel-brinda/masked-superstrings-supplement).

### 3.7 Experimental evaluation

#### Comparison of algorithms for superstring computation

We used KmerCamel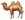 with *S. pneumoniae* as a model species, and considered for evaluations one assembled genome and one pan-genome (computed from 616 assemblies from a study of children in Massachusetts, USA [29]). To evaluate the behavior of the algorithms and representations for different types of de Bruijn graphs, we used varying the *k*-mer length to control the amount of branching along the whole genome, as well as shifting to the pan-genome to increase the amount of branching around polymorphic sites.

Using these data, we first sought to evaluate whether the proposed greedy heuristics can provide better superstrings than ProphAsm in those situations, in which ProphAsm (and more general all simplitigs) were unable to approach the lower bound given by the number of *k*-mers ([18, Fig 2]). The obtained results depict the number of superstring characters per distinct canonical *k*-mer (Fig. 2a); in the best case, the value would be close to 1.0, corresponding to nearly all positions in the masked superstring being unmasked. Note that the leftmost points, denoted by L1, correspond to local greedy extensions by one character, which is equivalent to the original ProphAsm algorithm, while the higher values correspond to higher maximal extension steps *d*_*max*_.

**Figure 2:**
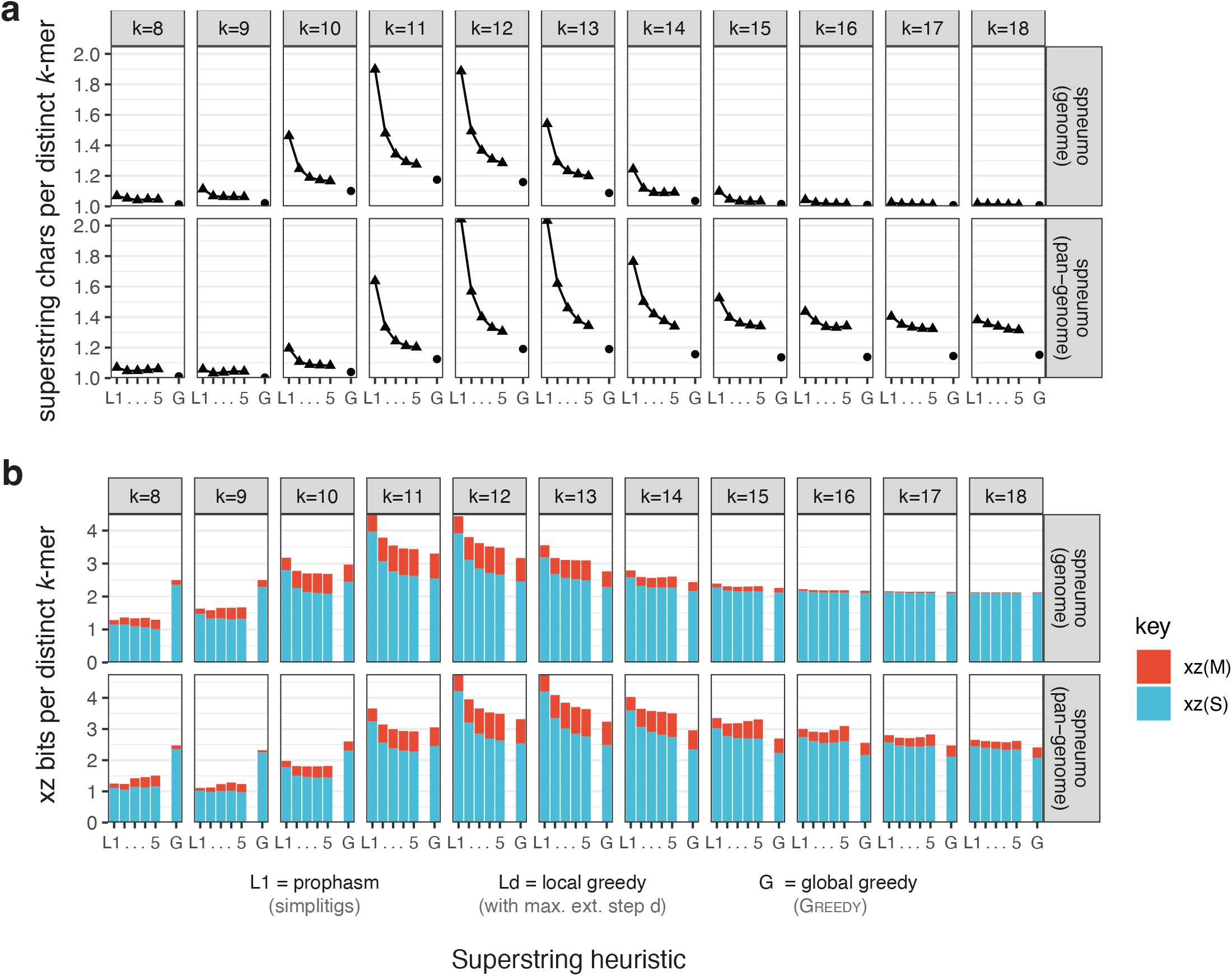
Comparison of greedy heuristics for *k*-mer superstrings. The benchmark was performed using a *S. pneumoniae* genome (NC_011900.1, *n* = 1, genome length 2.22 Mbp) and its pan-genome (dataset [29], *n* = 616, total genome length 1.27 Gbp), and evaluated for a range of *k*-mer lengths using BiDir-LocalGreedy (with *d*_*max*_ ∈ {1, …, 5}) and BiDir-GlobalGreedy, both in their hash-table-based implementations. **a)** Superstring (or mask) characters per distinct canonical *k*-mer; 1.0 is a (non-tight) lower bound. **b)** Bits per distinct canonical *k*-mer after superstring (*S*) and mask (*M*) compression (*‘*xz-9*’* of enc1 in Fig. 1, with the default masks produced by individual superstring heuristics).

We found that the global greedy heuristic always attained less than 1.2 characters per *k*-mer, providing a very good performance even for pan-genomes, irrespective of the specific value of *k*. For genomes, globally computed superstrings with larger values of *k* tend to attain nearly 1 character per *k*-mer, which is however not possible for pan-genomes, due to the higher interconnectivity of the graph around polymorphic sites. The local heuristic was nearly as successful in minimizing the superstring length as the global one, especially once a bigger maximal extension steps *d*_*max*_ was used, although a gap of up to 0.4 bits per *k*-mer was present even for higher values of *k*. However, in some situations, a further increasing of *d*_*max*_ led to worse results as longer extensions started blocking some potential long overlaps, that would the *k*-mer be able to form otherwise.

We further evaluated the compressibility of both the superstring and the mask (Fig. 1b). The experiments were performed with the default masks produced by individual superstring heuristics without further mask optimizations. Here, we observed two general modes of behavior. With *k* > log_4_ |*K*|, the *k*-mer sets were compressed to between 2 and 3 bits per *k*-mer, in dependence on the complexity of the corresponding de Bruijn graph. On the other hand, for *k* ≪ log_4_ |*K*|, xz could achieve compression below 2 bits per *k*-mer,attaining in extremal cases nearly 1 bit per *k*-mer, although only with local approaches and data structures producing regular patterns (i.e., hash tables, but not Aho-Corasick). This is the consequence of the fact that for extremely small values of *k*, nearly all possible *k*-mers are present, and the effective encoded information is the complement of the *k*-mer set. We also observed that in some situations, compressed superstrings might be improved by a higher *d*_*max*_, while simultaneously the compressibility decreased by a larger factor (pan-genome, *k* = 15).

To assess the generality of our finding, we sought to redo the analysis using bigger genomes, and used *S. cerevisiae* (n=1, genome length 12.2 Mbp) and a *SARS-CoV-2* pan-genome (n=14.7 M, total genome length 430 Gbp), and we found exactly the same patterns as previously with the pneumococcal data (data and plots provided in the supplementary Github repository https://github.com/karel-brinda/masked-superstrings-supplement).

#### Comparison of mask optimization algorithms

We envision the computation of masked superstrings as a two-step process, where the actual superstring is first computed with a pre-selected heuristic and accompanied by the default mask (dependent on the underlying data structures used for computing the superstring). Then, the default mask can be re-optimized for the specific end use-cases.

To evaluate the impact of mask optimization, we re-used the masked superstring computed previously and applied the following four re-optimization strategies: Default (D, preserving the current mask), MaxOne (O, maximizing the number of ones, i.e., unmasking all *k*-mers), MinOne (Z, minimizing the number of ones while greedily taking the leftmost ones), and MinRun (R, minimizing the number of runs).

We first evaluated the impact of individual heuristics on the compressibility (Fig. 3), finding several distinct modes of behavior. First, for extremely interconnected de Bruijn graphs, corresponding to *k* < log_4_ |*K*|, the MinOne and Default strategies provided the worst results; the reason is that most default ghost *k*-mers are in fact also in *K* and the attempts to avoid their re-occurrences only increased the entropy of the mask, whereas the other two strategies created contiguous, well-compressible segments. Second, for the local greedy algorithm with *d* = 1, producing simplitigs, the Default strategy provided the best compressible masks as they only contain runs of ones interrupted with (*k* − 1)-long stretches of zeros (as shown in Fig. 1c and Tab. 3), which is a highly compressible pattern. Third, for the other local and the global superstring heuristics, MinRun was always preferential and MaxOnes performed nearly equally well in most scenarios, while imposing much lower computational overhead.

**Figure 3:**
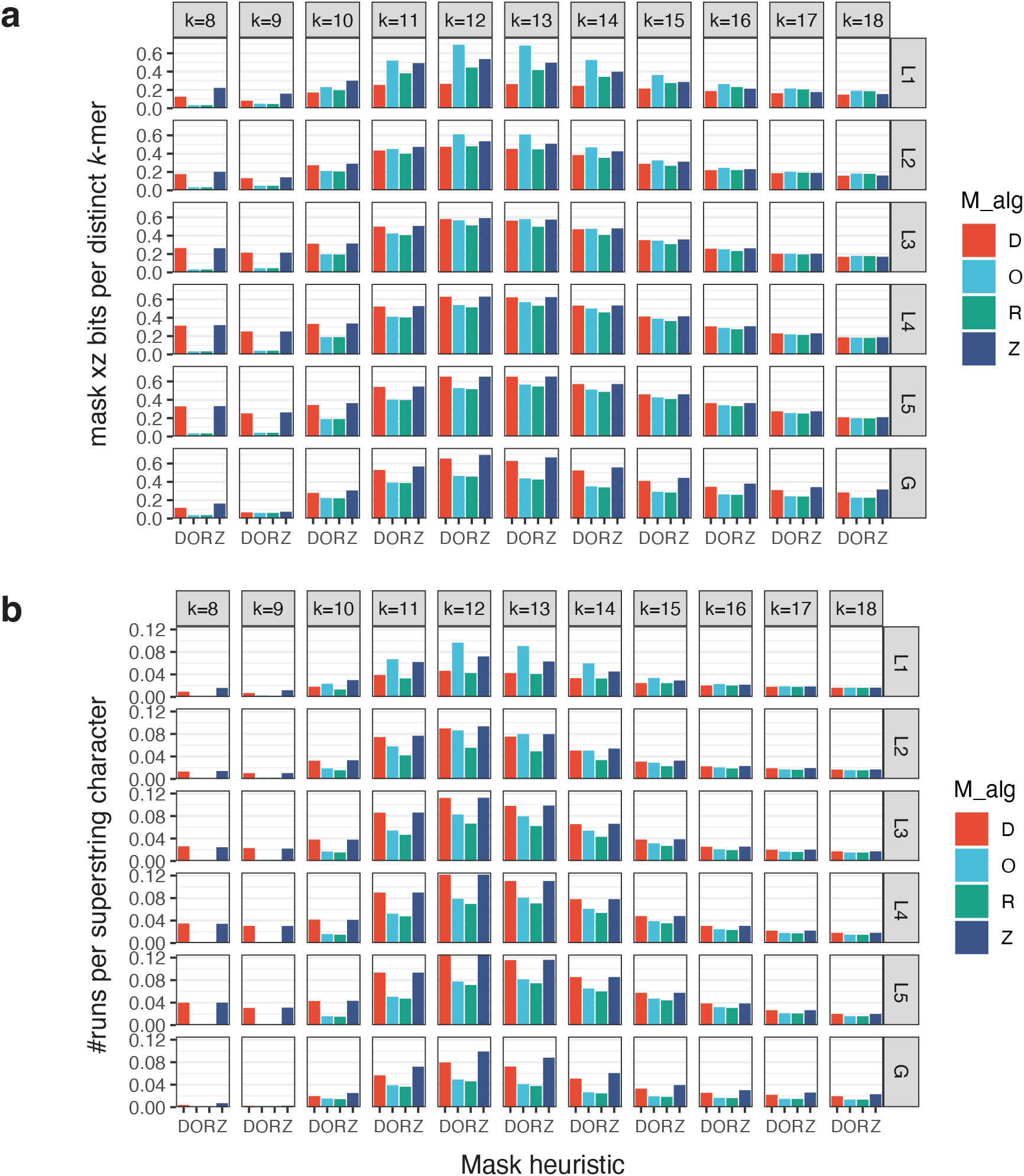
Comparison of four heuristics for mask optimization. The comparison was performed using the *k*-mer superstrings computed previously for the *S. pneumoniae* pan-genome. Individual heuristics: D – the default mask from the superstring algorithms, O – the maximal number of ones, Z – the minimum number of ones (computed greedily), and R – the minimal number of runs. **a)** Bits of mask after xz compression per distinct *k*-mer. **b)** Number of runs of ones per superstring character.

We then hypothesized that mask compressibility is proportional to the number of runs, and tested this using an analogical plot, depicting the number of runs per 1 superstring character (Fig. 3b). This number can be interpreted as the probability of starting a new block of *k*-mers within a superstring, and its lower values are better. We found that the observed patterns indeed clearly correlate with mask compressibility (Fig. 3a). Overall, this suggests that the best strategy is preserving default masks for the traditional (r)SPSS and minimizing the number of runs otherwise; if the latter is computationally infeasible, we suggest maximizing the number of ones.

#### Comparison to other existing approaches

Finally, we compared the masked superstring computed using the proposed local and global greedy heuristics to the state-of-the-art (r)SPSS representations, including eulertigs (optimal simplitigs) [21], and greedily and optimally computed matchtigs [22] (Tab. 1 and Tab. 2).

The results for the pneumococcal pan-genome in Tab. 1a and Tab. 2a imply that simplitigs (even optimal) and local greedy achieve worse results than global greedy or matchtigs. The performances of global greedy and matchtigs are comparable, with matchtigs being slightly better for the highly-branching case of *k* = 12. However, once we moved to datasets that do not satisfy the spectrum-like property that we created by subsampling the pneumococcal pan-genome to 10% of *k*-mers, we obtained a completely different story as displayed in Tab. 1b and Tab. 2b. Due to subsampling, the compressibility of the masked superstring worsened substantially for all algorithms, especially for local heuristics including simplitigs, matchtigs, and local greedy. The global greedy algorithm is now a clear winner, being more than two times better than matchtigs in terms of the final compressed sizes. Finally, we observe that local greedy*’*s performance improves with increasing *d*_*max*_.

## 4 Conclusions and discussion

In this paper, we developed a new concept for text-based *k*-mer set representations, called the masked superstring of *k*-mers, and showed it generalizes the existing (r)SPSS, comes with better compression properties, and provides additional flexibility for data that do not satisfy the spectrum-like property. We further demonstrated that the computation of optimal superstring and optimal masks are NP-hard tasks, we developed efficient local and global heuristic approaches, and implemented them in KmerCamel using hash tables and the Aho-Corasick automaton. We evaluated the proposed algorithms experimentally using viral, bacterial, and eukaryotic genomes and demonstrated that masked superstrings provide better compression characteristics than simplitigs, and can provide an improvement of up to several factors over the (r)SPSS, especially in applications involving subsampling.

**Table 1:**
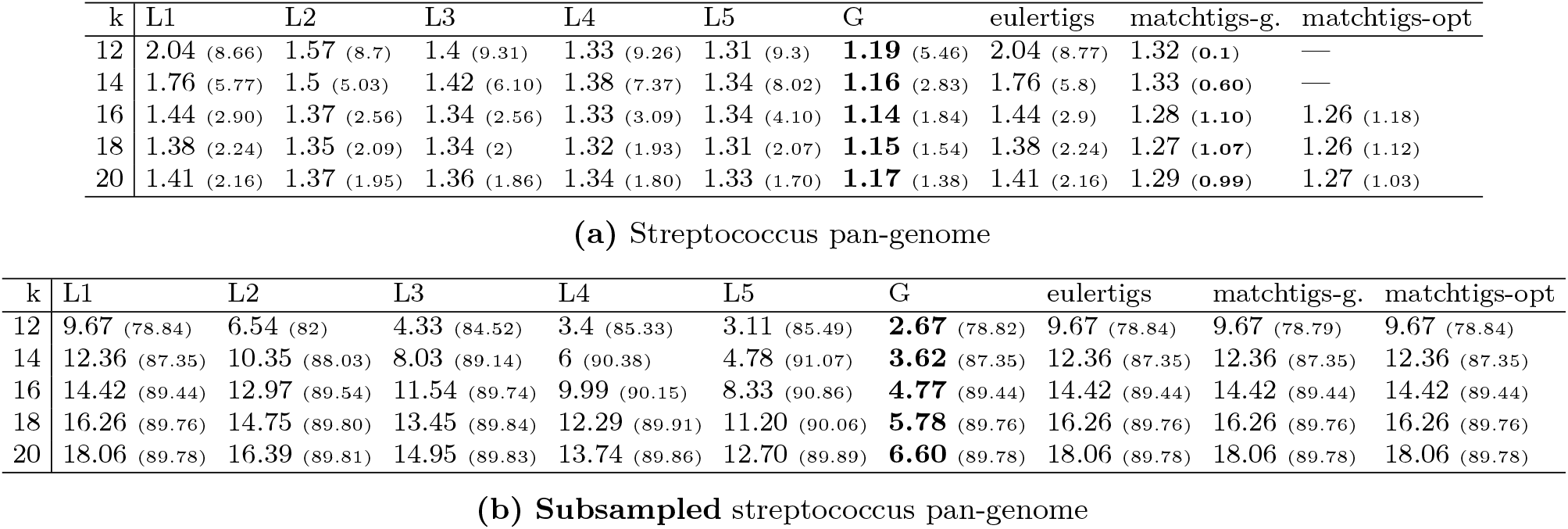
Comparison of mask characteristics. Comparison of our heuristics to (r)SPSS with respect to # of superstring characters *ℓ* (and with respect to 100 *⋅* # of runs *r* in *M*), both per distinct *k*-mer; G = global greedy, L*d* = local greedy (with maximum extension step *d*), and matchtigs-g. = greedily computed matchtigs. For all algorithms, the mask was optimized for the minimum number of runs. We were not able to compute optimal matchtigs for streptococcus pan-genome with *k* ∈ {12, 14} within reasonable time limit (several hours).

| k | L1 | L2 | L3 | L4 | L5 | G | eulertigs | matchtigs-g. | matchtigs-opt |
| --- | --- | --- | --- | --- | --- | --- | --- | --- | --- |
| 12 | 2.04 (8.66) | 1.57 (8.7) | 1.4 (9.31) | 1.33 (9.26) | 1.31 (9.3) | **1.19** (5.46) | 2.04 (8.77) | 1.32 (0.1) | — |
| 14 | 1.76 (5.77) | 1.5 (5.03) | 1.42 (6.10) | 1.38 (7.37) | 1.34 (8.02) | **1.16** (2.83) | 1.76 (5.8) | 1.33 (0.60) | — |
| 16 | 1.44 (2.90) | 1.37 (2.56) | 1.34 (2.56) | 1.33 (3.09) | 1.34 (4.10) | **1.14** (1.84) | 1.44 (2.9) | 1.28 (1.10) | 1.26 (1.18) |
| 18 | 1.38 (2.24) | 1.35 (2.09) | 1.34 (2) | 1.32 (1.93) | 1.31 (2.07) | **1.15** (1.54) | 1.38 (2.24) | 1.27 (1.07) | 1.26 (1.12) |
| 20 | 1.41 (2.16) | 1.37 (1.95) | 1.36 (1.86) | 1.34 (1.80) | 1.33 (1.70) | **1.17** (1.38) | 1.41 (2.16) | 1.29 (0.99) | 1.27 (1.03) |

(a) Streptococcus pan-genome
| k | L1 | L2 | L3 | L4 | L5 | G | eulertigs | matchtigs-g. | matchtigs-opt |
| --- | --- | --- | --- | --- | --- | --- | --- | --- | --- |
| 12 | 9.67 (78.84) | 6.54 (82) | 4.33 (84.52) | 3.4 (85.33) | 3.11 (85.49) | **2.67** (78.82) | 9.67 (78.84) | 9.67 (78.79) | 9.67 (78.84) |
| 14 | 12.36 (87.35) | 10.35 (88.03) | 8.03 (89.14) | 6 (90.38) | 4.78 (91.07) | **3.62** (87.35) | 12.36 (87.35) | 12.36 (87.35) | 12.36 (87.35) |
| 16 | 14.42 (89.44) | 12.97 (89.54) | 11.54 (89.74) | 9.99 (90.15) | 8.33 (90.86) | **4.77** (89.44) | 14.42 (89.44) | 14.42 (89.44) | 14.42 (89.44) |
| 18 | 16.26 (89.76) | 14.75 (89.80) | 13.45 (89.84) | 12.29 (89.91) | 11.20 (90.06) | **5.78** (89.76) | 16.26 (89.76) | 16.26 (89.76) | 16.26 (89.76) |
| 20 | 18.06 (89.78) | 16.39 (89.81) | 14.95 (89.83) | 13.74 (89.86) | 12.70 (89.89) | **6.60** (89.78) | 18.06 (89.78) | 18.06 (89.78) | 18.06 (89.78) |

(b) Subsampled streptococcus pan-genome

**Table 2:** Comparison of xz-compressed masked strings. Comparison of our heuristics to (r)SPSS with respect to # of xz bits of enc1 (*S* and *M*) per distinct *k*-mer; G = global greedy, L*d* = local greedy (with maximum extension step *d*). For all algorithms, the mask was optimized for the minimum number of runs. We were not able to compute optimal matchtigs for streptococcus pan-genome with *k* ∈ {12, 14} within reasonable time limit (several hours).

| k | # $k$-mers | L1 | L2 | L3 | L4 | L5 | G | eulertigs | matchtigs-greedy | matchtigs-opt |
| --- | --- | --- | --- | --- | --- | --- | --- | --- | --- | --- |
| 12 | 3336895 | 5.12 | 3.96 | 3.56 | 3.38 | 3.31 | 3.08 | 5.29 | **2.83** | — |
| 14 | 6089164 | 4.20 | 3.60 | 3.49 | 3.45 | 3.39 | **2.74** | 4.26 | 2.79 | — |
| 16 | 6945327 | 3.07 | 2.91 | 2.86 | 2.93 | 3.05 | **2.46** | 3.17 | 2.57 | 2.56 |
| 18 | 7367184 | 2.70 | 2.64 | 2.61 | 2.57 | 2.60 | **2.34** | 2.86 | 2.42 | 2.40 |
| 20 | 7731727 | 2.64 | 2.55 | 2.51 | 2.49 | 2.44 | **2.29** | 2.84 | 2.31 | 2.32 |

(a) Streptococcus pan-genome
| k | # $k$-mers | L1 | L2 | L3 | L4 | L5 | G | eulertigs | matchtigs-greedy | matchtigs-opt |
| --- | --- | --- | --- | --- | --- | --- | --- | --- | --- | --- |
| 12 | 333689 | 24.04 | 17.11 | 12.01 | 9.83 | 9.10 | **8.49** | 23.96 | 23.91 | 23.95 |
| 14 | 608916 | 28.35 | 24.51 | 19.91 | 15.64 | 12.96 | **10.84** | 28.24 | 28.24 | 28.24 |
| 16 | 694532 | 31.57 | 29.09 | 26.64 | 23.88 | 20.74 | **13.64** | 31.19 | 31.19 | 31.19 |
| 18 | 736718 | 35.04 | 32.48 | 30.31 | 28.37 | 26.49 | **16.00** | 34.43 | 34.43 | 34.43 |
| 20 | 773172 | 38.36 | 35.63 | 33.30 | 31.29 | 29.51 | **17.81** | 37.67 | 37.66 | 37.67 |

(b) Subsampled streptococcus pan-genome

Due to the experimental nature of our algorithms and their implementations, we focused in our evaluations predominantly on rather small genomes, which is the main limitation of this paper. However, we note that in many applications, such as bacterial pan-genomics, datasets tend to be small in terms of the overall *k*-mer diversity per one pan-genome. We also note that the proposed experimental methods, implemented in KmerCamel 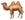, can be substantially optimized, especially the parts related to the Aho-Corasick automaton. Finally, the experiments with subsampled indexes demonstrate masked superstrings be suitable in applications where (r)SPSS could not be used efficiently.

Finally, multiple other challenges are to be addressed in order to transition masked superstrings to real bioinformatics applications. While the moving from unitigs to simplitigs/SPSS was nearly effortless in applications such as Bifrost [30] or SSHash [31], using masked superstrings in common data structures such as FM index [23] will require additional representation of the mask (while freeing other space). On the other hand, masked superstrings may enable a better compression in scenarios where the (r)SPSS representations were too limited, such as spaced-seed indexes in metagenomics [32].

Overall, we see masked superstrings as a unifying and generalizing concept that enables to better mathematically study and optimize textual *k*-mer-set representations, as well as it will provide new ways of increasing the efficiency bioinformatics software, especially in variation-heavy applications such as pangenomics.

## Acknowledgment

We are all grateful to Nicolaos Matsakis for the many fruitful discussions and for sharing his expertise on the shortest superstring problem. We also thank Ekaterina Milyutina for the discussions in the early stages of the project. O. Sladký and P. Veselý were partially supported by GA ČR project 22-22997S. P. Veselý was also partially supported by Center for Foundations of Modern Computer Science (Charles University project UNCE/SCI/004).

## A Overview of textual *k*-mer-set representations and the associated algorithms

**Table 3:**
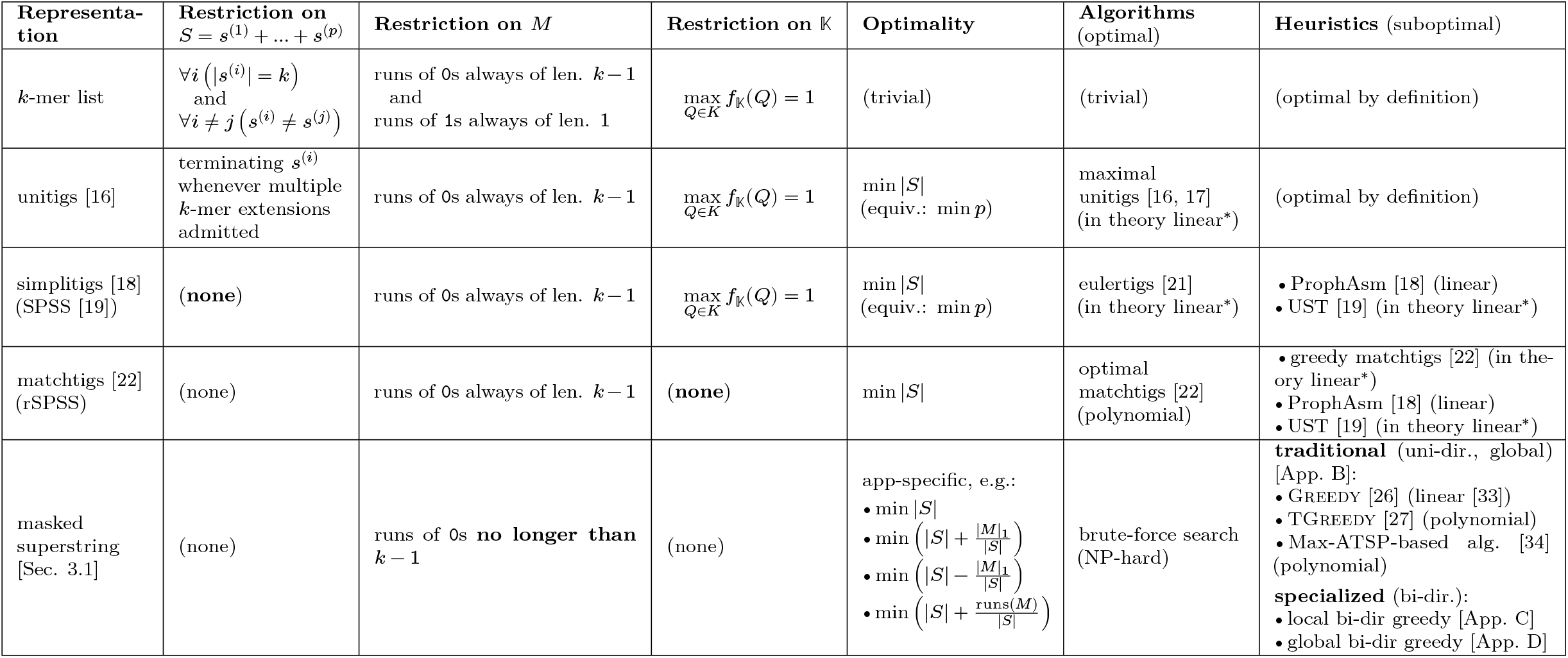
Overview of textual *k*-mer set representations and their constraints. The constraints are formulated over the superstring *𝒮* (and its components *s*^(1)^, …, *s*^(*p*)^ for (r)SPSS), the *k*-mer mask *M*, the multiset 𝕂 of all represented *k*-mers with their frequencies, the optimality criteria, the algorithms for computing the optimal representation, and the associated heuristics, including their time complexities. In bold, we highlight the change of the restrictions from the previous representation. _*∗*_In theory linear: The involved algorithms could in theory be linear, however their implementations involve the use of BCALM2 for computing unitigs, and as such do not have the guarantee of linearity. Remark about restrictions on 𝕏: None of these representations restricts the multiset 𝕏 of all ghost *k*-mers in any way (for representations with more components, ghost *k*-mers appear only if those components are concatenated into a superstring). We leave a study of restrictions on 𝕏 to a future work.

## B The shortest superstring problem

The *shortest (common) superstring problem* (SSP) is a fundamental and well-studied problem in the area of string algorithms, defined as follows: The input is a set of strings *𝒮* = {*s*_1_, *s*_2_, …, *s*_*m*_} over a finite alphabet Σ, and the goal is to compute a single string *𝒮* of minimum length such that *𝒮* contains every *s*_*i*_ as a substring. This *𝒮* is called the *shortest superstring* of *S*. In this appendix, we sketch algorithms developed for SSP as well as its main variants.

### Approximation algorithms and (global) greedy

SSP is NP-hard even when Σ is binary [35]. A standard way of dealing with computational hardness (at least in theory) is relaxing the requirement for the computed solution being optimal and designing approximation algorithms. We say that a polynomial-time algorithm *A* is *α-approximation* if the superstring computed by *A* has length at most *α* times the length of the optimal superstring. Despite more than three decades of study, the best possible approximation ratio for SSP is still widely open: The best known guarantee is *≈* 2.475 [36], while it has only been proven that computing a *≈* 1.003-approximation in polynomial time would imply P=NP [37].

One of the most important (and simplest) heuristics is the global greedy algorithm (called Greedy in algorithms literature), whose approximation ratio is between 2 and *≈* 3.425 [36] and which works as follows: Find two strings in *𝒮* that *overlap* the most (breaking ties arbitrarily), replace them in *𝒮* by their *merge*, and repeat until a single string remains in *S*, which is a superstring of the input strings. (For an example of overlaps, consider strings *𝒮* =*‘*ACGTGTGT*’* and *t* =*‘*TGTGTAA*’*, for which their largest overlap is ov(*s, t*) =*‘*TGTGT*’*, while ov(*t, s*) =*‘*A*’*, and merging string *s* to *t* (in this order) results in string *‘*ACG**TGTGT**AA*’*, with the overlap in bold.)

We describe global greedy in more detail in Appendix D, where we also develop its variant for *k*-mers superstring in the bi-directional model and describe an efficient (linear-time) implementation of this variant. In Appendix E, we provide another linear-time implementation, using the Aho-Corasick automaton.

### Overlap graph and cycle covers

There is a natural weighted directed graph *G*_*𝒮*_, called the *overlap graph*, associated with input *𝒮*: Each input string in *𝒮* is represented by a node and there is a directed edge from between any ordered pair of nodes (*s, t*) with weight equal to |ov(*s, t*)|, i.e., the length of the overlap when merging *s* to *t* (more precisely, the strings represented by *𝒮* and *t*). (This graph includes self-loops, with weight equal to the longest non-trivial self-overlap; for example, for *𝒮* =*‘*ACGTGTACG*’* we have ov(*s, s*) =*‘*ACG*’*.) We remark that, when *𝒮* is a set of *k*-mers, the vertex-centric de Bruijn graph of these *k*-mers is a subgraph of *G*_*𝒮*_ with edges corresponding to (*k* − 1)-long overlaps only.

Observe that any optimal solution for SSP corresponds to a longest Hamiltonian path in *G*_*𝒮*_ (i.e., a directed path that visits every node exactly once), however, computing such a path is, in general, NP-hard. It is nevertheless possible to efficiently compute an optimal *cycle cover*, which is a collection of directed cycles containing each node once and which may only have a larger total overlap than the optimal Hamiltonian path. A natural greedy algorithm outputs an optimal cycle cover, which can also be computed in linear time [38]. Note that cycle covers may be used for representing a set of strings as well, however, we leave their application for representing *k*-mer sets and an extension to the bi-directional model to a future work.

Overlap graphs and optimal cycle covers are useful when developing or analyzing an algorithm for SSP. In particular, they are central in the design of algorithms for which we can currently obtain substantially better approximation guarantees than for Greedy, such as TGreedy [27] or an algorithm based on the Max-ATSP problem [34, 36]. However, it is not known whether these two algorithms (or indeed any other algorithm) bring any real advantage compared to Greedy.

Lastly, we mention a variant of the overlap graph, called the *hierarchical overlap graph* (HOG) [39], which encodes maximal pairwise overlaps as well, but its size is linear in the input size (compared to quadratic for the overlap graph). Furthermore, linear-time constructions of HOGs have recently been designed [40, 41].

### Special case of same length strings

A special case of SSP, highly related to representing *k*-mers, is the *k*-SSP problem, in which all input strings have the same length *k*, though they may be over alphabets with possibly many letters. Solving *k*-SSP is NP-hard for any *k* ≥ 3, and in fact, hard to approximate arbitrarily well [42]. There is a handful of positive results: For *k* = 3 and 4, it is known that Greedy computes 2-approximate superstrings [43, 44, 45], while an algorithm by Golovnev et al. [46] gives better approximation guarantees than for general SSP when *k* ≤ 7.

### Other variants of SSP

Among the many variants of SSP in the literature, we only mention a few that seem most related to our application to *k*-mers. The *orientation-free SSP*, in which the superstring must contain each input string either as substring or a reverse of a substring, has been studied from the viewpoint of approximation algorithms [47, 48]; this variant may be seen as a step towards the bi-directional model of representing *k*-mers (considering just reverse strings instead of reverse complements). Cazaux and Rivals [49] propose a variant of SSP in which the superstring can be *circular* (i.e., it is written on a cycle and does not have a start or an end) and prove its NP-completeness.

## C Local greedy algorithm (generalized ProphAsm)

We develop a novel algorithm, called local greedy, for computing a masked superstring for a *k*-mer set. This algorithm generalizes ProphAsm [18], which compute heuristic simplitigs. In a nutshell, ProphAsm works by starting with an arbitrary *k*-mer *a* and finding a maximal path *P* in the (vertex-centric) de Bruijn graph that contains a (unlike for unitigs, this path may contain branching nodes); the *k*-mers covered by *P* are removed from the set. The process of finding a path is repeated until we obtain a collection of vertex disjoint paths that covers all input *k*-mers. Note that the path only contains edges corresponding to *k*-mer overlaps of length *k* − 1 (in the bi-directional model).

Local greedy generalizes these approaches by allowing to use overlaps shorter than *k* − 1, but not too much shorter. Local greedy has a parameter *d*_max_, the maximum depth, that specifies that all overlaps used by local greedy should be at least *k* − *d*_max_ long, i.e., we extend the superstring segment (corresponding to the current path) by at most *d*_max_ characters in each step.

In the uni-directional model, local greedy thus works as follows: Initially, let *K* be the set of *k*-mers. Until *K* gets empty, we take an arbitrary *k*-mer a ∈ K, remove it from K, and initialize a superstring segment *S*_*P*_ = a (that will correspond to a path in the overlap graph). We then repeatedly find a left or right extension of *S*_*P*_ which has maximum overlap with *S*_*P*_. A *left extension* of *S*_*P*_ by *d* characters (i.e., with overlap *k* − *d*) is a *k*-mer *b* from *K* such that *b* overlaps with a prefix of *S*_*P*_ by *k* − *d* characters; this amounts to finding *d* characters *a*_1_ …*a*_*d*_ such that *a*_1_ …*a*_*d*_ plus the first *k* − *d* characters of *S*_*P*_ belong to K. A *right extension* of superstring segment *S*_*P*_ is defined analogously, by extending the suffix of *S*_*P*_ by *d* characters.

In each step of extending *S*_*P*_, we find a left or right extension with the minimum value of *d* (i.e., with the longest overlap with *S*_*P*_). If *d* ≤ *d*_max_, then we append the *d* characters of the extension to *S*_*P*_ and the corresponding *k*-mer *b* to *P* (for left or right), and we also remove *b* from K. Otherwise, any left or right extension requires adding more than *d*_max_ characters, and we close *S*_*P*_, append *S*_*P*_ to the constructed superstring, and initialize new segment *S*_*P*_ if *K* is still non-empty. Finally, to extend local greedy into the bi-directional model, we take *K* as the set of canonical *k*-mers and instead of checking whether a *k*-mer *b* is in K, we test if the canonical *k*-mer of *b* is in *K* (recall that the canonical *k*-mer of a is the lexicographic minimum of a and its reverse complement).

We provide a pseudocode of local greedy in the bi-directional model, called BiDir-LocalGreedy, in Algorithm 1, where we implement searching for a left or right extension in a rather straightforward way by exhaustively enumerating all possible length-*d* strings and checking whether this gives a valid extension for the current path. This enumeration takes time 4^*d*^, but as we first start with *d* = 1 and keep increasing it by one only if no length-*d* extension is found, we do not get to a large value of *d* in many steps. Nevertheless, searching for an extension can take time up to 4^*d*^max, which is tolerable for a small value of *d*_max_, but can affect the running time dramatically if the current path has no short extension.

### Alternative implementation

In Appendix E, we describe an implementation of local greedy using the Aho-Corasick automaton that avoids this exhaustive enumaration and runs in linear time (in the total length of all *k*-mers). This automaton-based implementation outperforms the straightforward (but exponential-time) implementation in Algorithm 1 for large values of *d*_max_. Nevertheless, using a tree-based data structure leads to certain overheads (such as much higher memory usage or worse memory access patterns) that make the straightforward implementation preferable for small-enough values of *d*.

#### Algorithm 1

BiDir-LocalGreedy – outputs a superstring of *K* and its mask. The case *d*_*max*_ = 1 corresponds to Alg. 1 in [18] (ProphAsm).

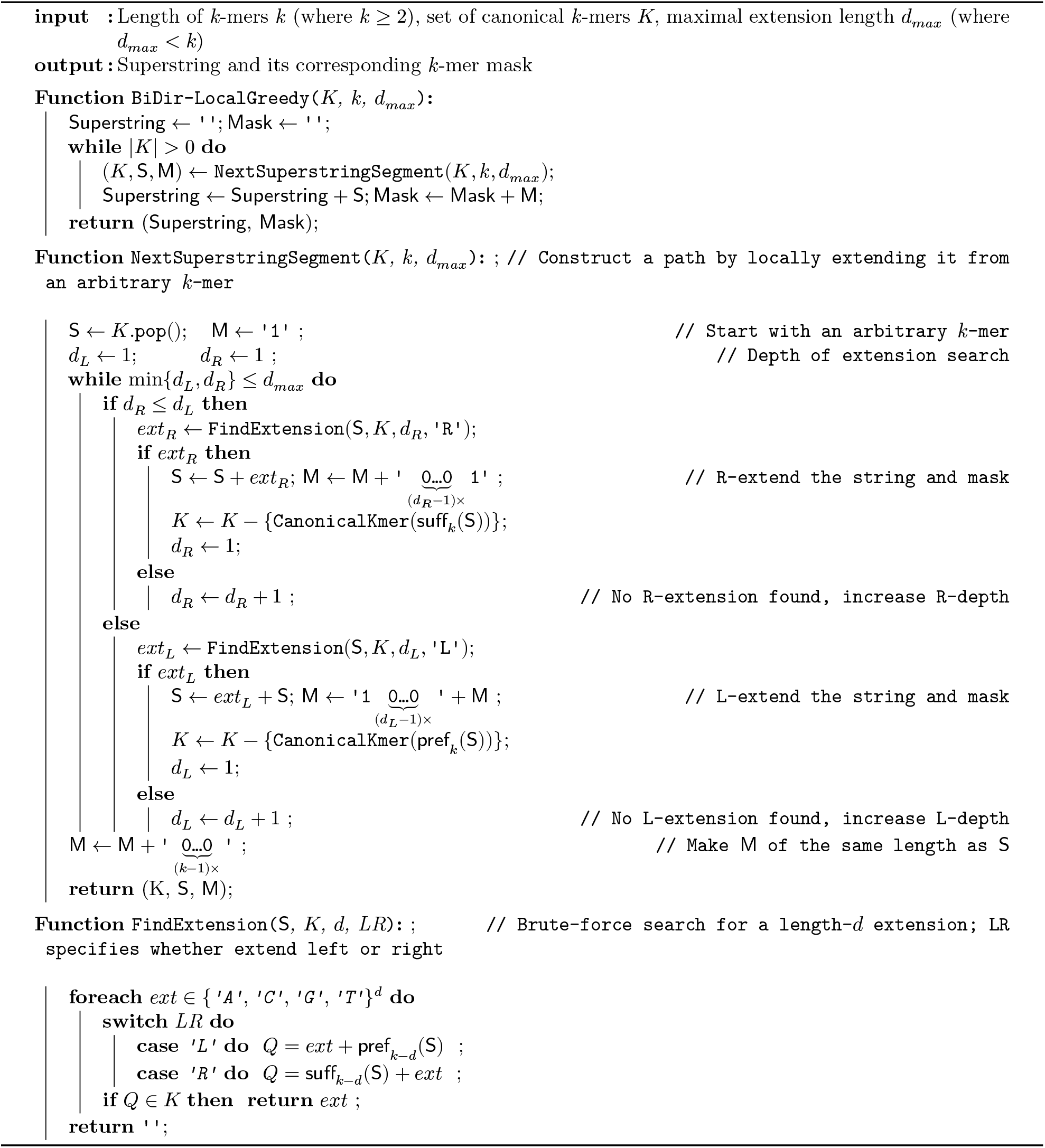

## D Global greedy algorithm in the bi-directional model (BiDir-GlobalGreedy)

In this appendix, we describe the global greedy algorithm for finding superstrings and develop its adjustment for representing *k*-mers in the bi-directional model, together with an efficient hashing-based implementation.

### Global greedy algorithm for the shortest superstring problem

The **global greedy algorithm**, called Greedy in the algorithms literature, is one of the most important (and simplest) heuristics for the Shortest Superstring Problem (SSP); cf. [26, 27, 36] (see Appendix B for an overview of SSP). Greedy works as follows: Find two strings in the set of input strings *𝒮* that *overlap* the most (breaking ties arbitrarily), replace them in *𝒮* by their *merge*, and repeat until a single string remains in *S*, which is a superstring of the input strings. (For an example of overlaps, consider strings *s* =*‘*AGATA*’* and *t* =*‘*TAGCC*’*, for which their overlap ov(*s, t*) =*‘*TA*’*, while ov(*t, s*) is the empty string, and merging string *s* to *t* (in this order) results in string *‘*AGA**TA**GCC*’*, with the overlap in bold.) See Figure 4 for an example.

**Figure 4:**
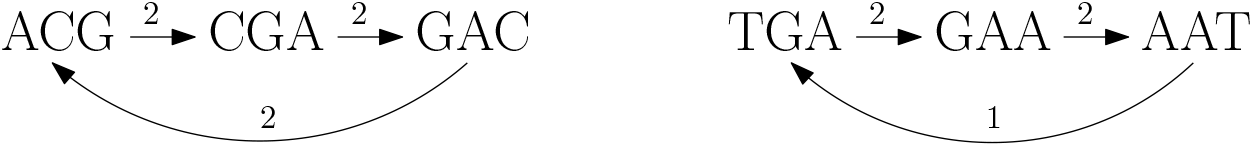
The optimal superstring in this simple example (in the uni-directional model) is *‘*ACGAATGAC*’* with length 9. On the other hand, Greedy will first construct string *‘*ACGAC*’* which will later concatenate to the constructed string *‘*TGAAT*’*, giving length of 10. All overlaps for Greedy consist of two characters, except for the concatenation, which consists of 0 characters. The curved edges cannot be part of the Greedy output.

In terms of the overlap graph *G*_*𝒮*_ of input strings *𝒮*, Greedy builds a Hamiltonian path *H* in *G*_*𝒮*_ by going over all of the edges in *G*_*𝒮*_ in the order of non-increasing overlap lengths (breaking ties arbitrarily; in the example above, edge (*s, t*) has overlap length of 5). It adds the currently processed edge (*s, t*) to *H* if *s* has outdegree 0 in *H, t* has indegree 0 in *H*, and edge (*s, t*) does not close a cycle in *H*. Since *G*_*𝒮*_ is complete directed graph, *H* is indeed a Hamiltonian path after we process all edges.

Greedy has been a popular choice for computing superstrings in practice [50, 51, 52], thanks to its simplicity, good practical performance [53, 54], and linear-time implementations [33], which we describe in Appendix E. Somewhat surprisingly, the worst-case performance of Greedy in terms of the superstring length is not known yet. The best upper bound on its approximation ratio is *≈* 3.425 [36], while there are simple examples showing that Greedy may output a superstring of length about two times the length of the optimal superstring. In fact, Tarhio and Ukkonen [43] conjectured already in the 80s that its approximation ratio is 2; this conjecture is still open.

Nevertheless, Greedy provides 1/2-approximation for the compression measure [43, 55], which quantifies how many input characters can be saved from the raw input (i.e., it equals the total length of input strings minus the length of the superstring); this ratio of 1/2 is tight for Greedy.

### BiDir-GlobalGreedy for representing *k*-mer sets in the bi-directional model

First, we observe that in the uni-directional model, we can use this greedy algorithm directly to get a superstring for a given *k*-mer set. Indeed, if we do not take reverse complements into account, we just aim to find a superstring for an input set of strings, which are the individual *k*-mers.

We now modify the global greedy algorithm for the bi-directional model so that it utilizes the possibility to merge a *k*-mer with a reverse complement (RC) of another *k*-mer and to represent either the original *k*-mer or its RC but not necessarily both. In other words, a naïve application of global greedy in the bi-directional model may result in unnecessarily long superstrings. We then show that the adjusted algorithm can still be implemented in linear time.

For each *k*-mer present in the *k*-mer set, we consider both the *k*-mer and its RC, and we forbid using an edge between them (forbidding edges between a *k*-mer and its RC allows us to maintain a certain useful invariant as we show below). Similarly as in the uni-directional model, our aim is to greedily construct a Hamiltonian path *H* in the overlap graph. However, in the bi-directional model we ensure that no *k*-mer and its RC both appear in *H*; the aim is to make the resulting superstring possibly shorter. In other words, we adjust the Hamiltonian requirement so that the resulting path *H* contains each *k*-mer or its RC but not both. (While we restrict *H* in this way for efficiency reasons, a similar property does not hold for the resulting superstring *S* obtained by merging *k*-mers in *H* in the order given by *H*. Indeed, a *k*-mer a in *H* may appear more times in *S* or the RC of a may be a substring of *S* as a result of merging some other *k*-mers.)

We therefore modify the global greedy algorithm as follows:

- We maintain a set of chosen edges *H* in the overlap graph for the *k*-mer set, where each *k*-mer also has its RC.
- In each step, we choose a largest-overlap edge from *k*-mer a to *k*-mer *b* such that a has outdegree 0, *b* has indegree 0, the edge does not close a cycle, and *b* is not an RC of a (breaking ties arbitrarily). Letting a^′^ denote the RC of a and *b*^′^ the RC of *b*, we add edges (a, *b*) and (*b*^′^, a^′^) to *H*.

Alternatively, BiDir-GlobalGreedy can be described by merging strings: In each step *t*, there is a set *S*_*t*_ of strings to merge (initially, this is the set of *k*-mers and their RCs). In *𝒮*_*t*_, choose two different strings a and *b* such that their overlap is the longest (when merging a to *b* in this order) and *b* is not an RC of *a*, breaking ties arbitrarily. Letting *a*^′^ denote the RC of *a* and *b*^′^ the RC of *b*, we merge a to *b* and also merge *b*^′^ to *a*^′^.

This way, BiDir-GlobalGreedy maintains the following invariant: *In every step t, it holds that for each string s ∈ S*_*t*_, *the RC of s is also present in S*_*t*_ (formally, this can be shown by mathematical induction). This invariant in particular implies that in each step, *b*^′^ has outdegree 0 and a^′^ has indegree 0.

While global greedy in the uni-directional model outputs a single Hamiltonian path, in the bi-directional model we end up with two disjoint paths of the same length, each satisfying the adjusted Hamiltonian property, which correspond to two reverse complementary superstrings; this follows directly from the aforementioned invariant.

The linear-time implementation of Greedy using the Aho-Corasick automaton [33] can be extended to handle our modification; see Appendix E. However, in case of *k*-mers, the automaton is not needed for linear-time complexity of *𝒪*(*k ⋅ n*), where *n* is the number of *k*-mers. Moreover, a simpler hashing-based implementation is often preferable in terms of the running time or memory usage; its main downside is that it works for *k* < 32 (when we use 64-bit integers to store *k*-mers), while the automaton-based implementation does not restrict *k*.

### Hashing-based implementation of BiDir-GlobalGreedy

For each overlap length *d* from *k* − 1 to 0, we create a hash table mapping each existing prefix of size *d* to a list of *k*-mers with this prefix which so far have indegree 0. Then, for each *k*-mer a with outdegree 0, we find the first *k*-mer with length-*d* prefix equal to the length-*d* suffix of a such that:

- The corresponding directed edge does not form a cycle, which we check similarly as in [33]. In particular, as *H* is a collection of paths during the computation, for each path *P* in *H* we maintain pointers between the endpoints of *P* (these are arrays first and last in the pseudocode).
- The edge does not go to a *k*-mer with indegree 1 (although we filter out nodes of non-zero indegree when creating the hash table, the indegrees may have changed as we are adding edges between reverse complementary *k*-mers). Whenever this happens, we erase this *k*-mer from the prefix hash table.
- The edge does not go from a string to its reverse complement.

We provide a pseudocode for this hashing-based implementation of BiDir-GlobalGreedy in Algorithm 2.

The construction of the prefix hash table runs in expected time *𝒪*(*k ⋅ n*) provided that we can compute any prefix (or suffix) of a *k*-mer with bit operations in constant time; namely, our implementation using 64-bit integers thus requires for *k* < 32. Therefore, the only potentially expensive part is finding the first *k*-mer in the prefix table that fulfills the conditions above.

Note, however, that this does not increase the time complexity. For a fixed *d*, the second case can happen in total at most *n* times as we can remove each *k*-mer only once, and similarly, the first case can happen at most once per *k*-mer [33]. The same holds for the third case as for each string there is only one complementary string and this complementary string has only one *k*-mer with indegree 0.

Therefore, the algorithm runs in expected time *𝒪*(*k ⋅ n*), that is in linear time with respect to the total length of the *k*-mers. We can achieve this complexity in the worst case if we use the implementation using the Aho-Corasick automaton (which can also work with *k* ≥ 32), at the cost of a certain overhead of using a prefix-tree-based data structure; see Appendix E.

#### Algorithm 2

BiDir-GlobalGreedy-Hashing – a hashing-based implementation of the global greedy algorithm that computes a superstring representation of a set of *k*-mers *K* in the bi-directional model.

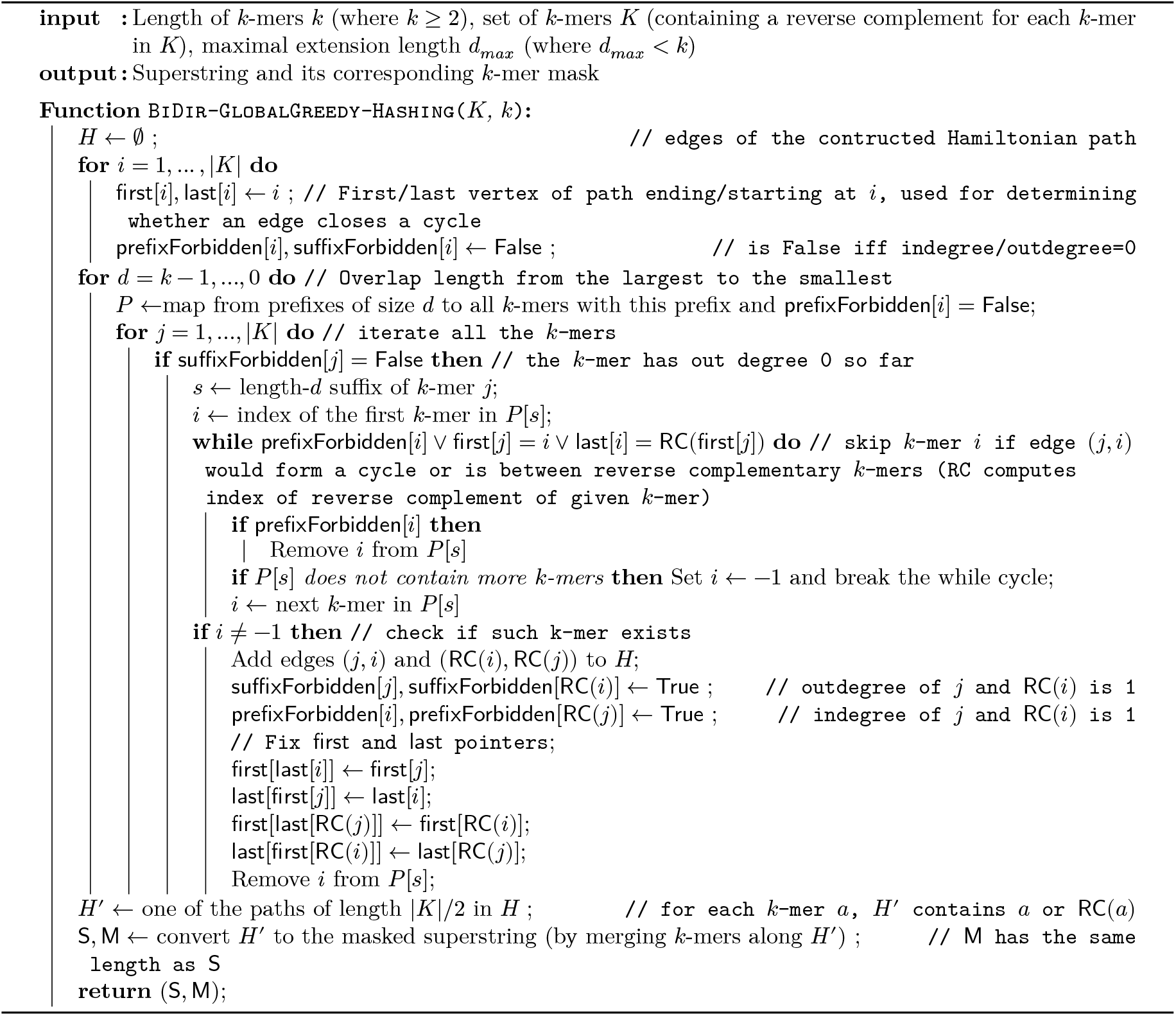

## E Implementations of local and global greedy using the Aho-Corasick automaton

In this appendix, we develop worst-case linear-time implementations for both local and global greedy algorithms using the Aho-Corasick (AC) automaton. For local greedy, this automaton-based implementation thus avoids an exhaustive enumeration, but introduces a certain overheads due to using a tree-based data structure.

A linear-time implementation of Greedy for SSP using the AC automaton was already designed by Ukkonen [33], and we extend it to representing *k*-mers in the bi-directional model, that is, for BiDir-GlobalGreedy, using similar ideas as in Appendix D. However, a hashing-based implementation of BiDir-GlobalGreedy also runs in (expected) linear time and does not have overheads caused by the automaton. Therefore, the hashing-based implementation should be preferable, and we leave it to a future work to develop a more efficient implementation of BiDir-GlobalGreedy.

### Aho-Corasick automaton

Formally, the Aho-Corasick (AC) automaton is a trie (prefix tree) constructed from the input string (in our case *k*-mers) equipped with a failure function *f*, where *f* (*s*) is the state of the longest proper suffix of state *s* that corresponds to a state of the trie (i.e., is a prefix of an input string). Abusing notation, we denote by *s* the string (which is a prefix of an input string) corresponding to state *s*. The automaton further has an output function which for a given state *s* iterates over all input strings that are suffixes of *s*. For representing *k*-mers, the output function is trivial as the states corresponding to *k*-mers are always leaves of the trie. The automaton can be constructed in linear time; we refer to a standard algorithms textbook (e.g. [56]) for a more detailed description. See Figure 5 for an illustration.

**Figure 5:**
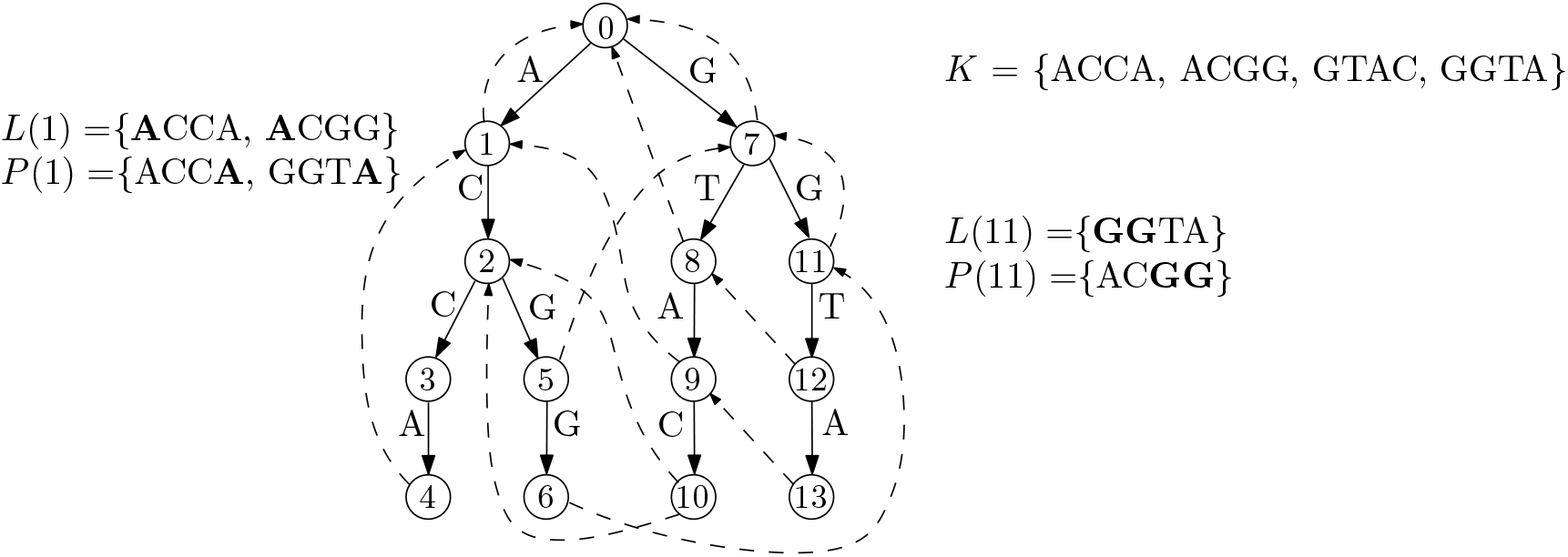
An illustration of the AC automaton constructed from *k*-mers ACCA, ACGG, GTAC, GGTA (with *k* = 4) in the uni-directional model. Forward edges (solid) are labeled by letters ACGT, while the failure function *f* is depicted by dashed edges (which are not part of the trie of the *k*-mers). State 0 corresponds to the empty prefix and is the root of the trie, while, for example, state 2 corresponds to prefix AC and state 9 to prefix GTA. We also provide examples of lists *L*(*s*) (with *k*-mers that have *s* as their prefix) and *P* (*s*) (with *k*-mers having *s* as their suffix) for *s* = 1 and 11; these lists are used in the implementation of global greedy. (Technically, lists *L*(*s*) and *P* (*s*) store just indices of *k*-mers, not the whole *k*-mers.)

The AC automaton has found many applications. Besides its standard use for string searching and the aforementioned implementation of Greedy for SSP in linear time [33], it was used to show a link between the Burrows-Wheeler Transform and the eXtended Burrows-Wheeler Transform; see e.g. [57].

### BiDir-GlobalGreedy using the AC automaton

We first describe how to use the automaton to implement global greedy for *k*-mer set representation in the uni-directional model; this is essentially the same implementation as in [33]. For every state *s* in the automaton, we maintain a list *L*(*s*) of *k*-mers which have *s* as a prefix, and another list *P* (*s*) for those which have *s* as a suffix. Then, we traverse the automaton in the reverse breadth first search (BFS) order with the aim to construct a Hamiltonian path *H* in the overlap graph. The order of traversal guarantees that the pairs of *k*-mers with the highest overlap are merged first.

When we visit a state *s*, we use lists *L*(*s*) and *P* (*s*) to find a pair (a, *b*) of different *k*-mers such that:

- a has *s* as its prefix and *b* has *s* as its suffix,
- edge (a, *b*) does not close a cycle in *H*
- a has outdegree 0 in *H* and *b* has indegree 0 in *H*,

Namely, for each a *∈ L*(*s*), if a has outdegree 0 in *H*, we iterate *b ∈ P* (*s*) and check the conditions above (that is, while visiting a state we may add more edges, at most one for each a *∈ L*(*s*)). Note that the first condition is ensured by taking a *∈ L*(*s*) and *b ∈ P* (*s*). The second condition may not be satisfied only once for each *k*-mer a and each of at most *k* − 1 states that contain it in *L*(*s*), that is, we reject an edge due to the second condition at most *𝒪*(*k ⋅ n*) times. For the last condition, we just need to check whether *b* has indegree 0 in *H* and if not, we remove *b* from *P* (*s*). This ensures that the implementation runs in linear time; see [33] for details. Finally, having a complete Hamiltonian path *H*, we merge *k*-mers in the order given by *H*. This concludes the description for the uni-direction model.

In the bi-directional model, we use a similar modification as for the hashing-based implementation described in Appendix D: When we add an edge (*a, b*) to *H*, we also add (*b*^′^, *a*^′^) to *H*, where *a*^′^ and *b*^′^ are reverse complements (RCs) of a and *b*, respectively. Further, we forbid using edges that lead between a *k*-mer and its RC. This way, we eventually construct two Hamiltonian paths.

### Local greedy using the AC automaton

We use the automaton to decrease the time complexity of finding the left/right extension in the local greedy algorithm, described in Appendix C. In the version with *k*-mer hashing, this has exponential-time complexity with respect to the current value of *d*_*L*_ or *d*_*R*_, which may be up to *d*_*max*_ in the worst case. (Recall that *d*_*max*_ is a parameter of local greedy, which specifies that the overlap in any step is at least *k* − *d*_*max*_).

As in global greedy, for each state *s* of the automaton, we maintain the lists *L*(*s*) and *P* (*s*) of *k*-mers which have the string corresponding to *s* as a prefix or a suffix, respectively. Furthermore, for each prefix and suffix of each *k*-mer we store the state of the automaton corresponding to the prefix/suffix (such state may not exist for the suffix).

Suppose we are looking for a left extension of length *d* and let *s* be the length-(*k* −*d*) prefix of the currently constructed string (denoted *S* in function NextPath in Algorithm 1). Since *s* is a length-(*k* − *d*) prefix of some *k*-mer *a*, state *s* is in the automaton and we iterate *P* (*s*) to find a *k*-mer not equal to a or its RC which has not been used as an extension or a starting *k*-mer yet. To ensure that this runs in linear time, when we find a *k*-mer *b* in *P* (*s*) that has been used already, we remove *b* from *P* (*s*).

We find a right extension analogously, with *s* being the length-(*k* − *d*) suffix of the currently constructed string *S*. If *s* is not a state of the automaton, there is no length-*d* extension, and otherwise, we iterate list *L*(*s*) similarly as we loop over *P* (*s*).

We argue that this implementation runs in time *𝒪*(*k ⋅ n*) (that is, linear in the total length of the *k*-mers). The automaton and the mapping from prefixes and suffixes of *k*-mers to automaton states can be constructed in linear time. Using this mapping, searching for a left or right extension of length *d* can be implemented in *𝒪*(1) time if we do not count removals from lists *L*(*s*) and *P* (*s*). Since every *k*-mer appears in *k* − 1 lists *L*(*s*) and *k* − 1 lists *P* (*s*), there are at most *𝒪*(*k ⋅ n*) removals from these lists (each taking *𝒪*(1) time).

### Using other data structures for strings

We implemented the local and global greedy algorithms in worst-case linear time using the AC automaton, which is a tree-based data structure and thus, may have some overheads compared to, say, hash tables. We leave it to a future work whether these algorithms can be implemented in worst-case linear time using more efficient data structures for strings, such as the FM-index [23].

## F NP-hardness of finding the shortest superstring of *k*-mers

In this appendix, we show NP-hardness of finding a minimum-length superstring of a *k*-mer set, but we do not consider masks for this superstring and their compressibility (complexity of and heuristics for mask optimization are discussed in Appendices G and H).

### Theorem F.1.

*Given a set of k-mers K, with k* = Θ(log |*K*|), *finding the shortest superstring for K is NP-hard in both the uni-directional and bi-directional models*.

We will show that the theorem follows from a corresponding hardness of the shortest superstring problem (SSP), in which we need to find the minimum-length string *S* that contains every input string as substring. SSP is well-known to be NP-hard even for a binary alphabet [35] (in fact, it is NP-complete as its decision version belongs to NP). However, this does not immediately imply that SSP is NP-hard when input strings are *k*-mers or in the bi-directional model, where we allow to merge a *k*-mer with a reverse complement (RC) of another *k*-mer and each *k*-mer may be represented in *S* either as a substring or an RC of a substring.

Considering *k*-mers (in the uni-directional model) basically corresponds to requiring that all input strings for SSP have the same length *k* and are from an alphabet with at most four letters, which can be renamed to {A, C, G, T}. It turns out that this special case of SSP is still NP-hard, in fact even for a binary alphabet:

### Proposition F.2

(Theorem 3 in [26]). *The shortest superstring problem is NP-hard even if the input strings are over the binary alphabet and have the same length k, for any k* = Ω(log *n*), *where n is the number of input strings*.

*Proof of Theorem F*.*1*. Given an SSP input satisfying restrictions of Proposition F.2, we transform it into alphabet {A, C} (changing all 0s to As and all 1s to Cs), which defines the input *k*-mer set K. This shows NP-hardness for the uni-directional model. To see that the hardness holds even in the bi-directional model, observe that the reverse complement of any *k*-mer from *K* belongs to {G, T}_*k*_ and thus, there is no non-trivial overlap between a *k*-mer in *K* and an RC of a *k*-mer in K. Hence, considering reverse complements does not bring any advantage when optimizing the superstring length.

In more detail, we claim that there is an optimal superstring not containing letters G and T (thus consisting of As and Cs only); such a superstring does not contain the RC of any original *k*-mer as substring. Indeed, let *S* be any superstring such that every *k*-mer in *K* or its reverse complement appears as a substring in *S*. We modify *S* into a string *S*^′^ by taking every maximal interval consisting of characters {G, T} only and replace it with its RC. This operation does not change the length and ensures that *S*^′^ still represents all *k*-mers, since any maximal interval with letters G and T only has no partial overlap with any *k*-mer or its RC. This shows the claim.

*Remark* F.3. Note that while small values of *k* (say *k* ≤ 32) are often sufficient in practice, the NP-hardness proof requires an arbitrarily large *k*. In fact, it is not possible to show a hardness result for a fixed *k* — indeed, since the alphabet size is four, there are at most 4^*k*^ possible *k*-mers, so the problem would have a constant size for a constant *k*. Thus, the hardness result is tight up to constant factors as it only requires *k* = Ω(log |*K*|).

## G NP-hardness of minimizing the number of runs in a mask

We show that for a given superstring finding a mask that minimizes the number of runs of ones is NP-hard (this optimization can be thought of as improving compressibility, namely, it minimizes the run length encoding). We show the hardness in the uni-directional model, where we do not consider *k*-mer and its reverse complement as equivalent. Nevertheless, we conjecture that the hardness holds in the bi-directional model as well. More formally, we consider the following problem, called MaskMinNumRuns: Given a set of *k*-mers *K* (for an arbitrary *k* ≥ 2), and their superstring *S*, find a mask for *S* (w.r.t. K) that has the minimum number of runs of ones. Recall that a binary string *M* of the same length as *S* is a *mask* for *S* (w.r.t. K) if every *k*-mer a ∈ *K* has at least one occurrence in *S* masked with 1 (that is, there is *i* such that *M*[*i*] = 1 and *S*[*i* ∶ *i* + *k*] = *a*, where *S*[*i* ∶ *i* + *k*] is the length-*k* substring of *S* starting at *i*) and for every index *i*, if *S*[*i* ∶ *i* + *k*] is not a *k*-mer in K, then *M*[*i*] = 0. We prove the following theorem:

### Theorem G.1.

*MaskMinNumRuns in the uni-directional model is NP-complete. Furthermore, the problem is NP-hard to o*(log |*K*|)*-approximate, i*.*e*., *it is NP-hard to find a mask with o*(log |*K*|) *times the optimal number of runs of ones*.

Note that *k* must not be bounded; this follows from a similar reason as outlined in Remark F.3. As the problem is clearly in NP (with mask being a certificate, whose validity can be verified in polynomial time), it is sufficient to show NP-hardness. Strictly speaking, we prove the NP-hardness for the decision version of MaskMinNumRuns, which asks to determine whether there is a mask with at most *ℓ* runs of ones, for a given *ℓ*.

The approximation hardness uses the following (tight) result about Set Cover:

### Theorem G.2

(Corollary 4 in [58]). *For every fixed* ϵ > 0, *it is NP-hard to approximate Set Cover to within an* (1 − ϵ) *⋅* ln *N factor, where N is the size of the instance*.

*Proof of Theorem G*.*1*. First, we restate the MaskMinNumRuns problem using graph theory. We can take the edge-centric de Bruijn graph of the superstring and color the edges with two colors – blue if it corresponds to a ghost *k*-mer, i.e., a length-*k* substring not in *K*, and red if it appears in K. We can now observe that the superstring corresponds to a walk *W* in the de Bruijn graph. We can reformulate our problem as selecting the smallest number of subwalks of *W* consisting only of red edges such that all red edges are covered by one of the selected subwalks. Observe that whenever we add a red edge into a selected subwalk, we can select all succeeding and preceding red edges in *W* until we reach a blue edge in *W*, without having to add another subwalk. Therefore, we can split *W* by blue edges into maximal red subwalks and find the minimum number of these red subwalks we need to take in order to cover all red edges in the graph.

With this formulation in hand, we show a reduction from Set Cover. Recall that an instance of Set Cover consists of universe *U* and set of *m* subsets *A*_1_, …, *A*_*m*_ of *U*, and the goal is to select the minimum number of these subsets *A*_*i*_ that cover *U*, i.e., whose union equals *U*.

Given an instance of Set Cover, we choose *n* ≥ max{|*U*|/2, 4} and take a complete de Bruijn graph *G* with *n* vertices corresponding to all strings of length *k* − 1 = log_4_ *n* over the ACGT alphabet. The *k*-mers of our instance (including ghost *k*-mers) will correspond to a subset of the 4*n* edges of *G*, with each edge representing the length-*k* merge of its two endpoints. Note that *G* has two edge-disjoint Hamiltonian cycles which we denote *H*_*B*_ and *H*_*R*_ and which can be found in polynomial time [59] (in fact, there are exactly three edge-disjoint Hamiltonian cycles).

We color edges of *H*_*B*_ in blue and the edges of *H*_*R*_ in dark red. We color all the remaining edges (not in *H*_*B*_ or *H*_*R*_) in light red. As *G* contains 4*n* edges in total, out of which we colored *n* in blue and *n* in dark red, there are 2*n* light-red edges. We map each element in *U* to a light-red edge and delete the unmapped light-red edges. This way, we get a bijection between light-red edges and *U*. Furthermore, we modify every set *A*_*i*_ by adding all the dark-red edges to obtain new sets denoted 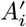. We also add (new elements corresponding to) the dark-red edges into *U* and a modified universe *U* ^′^. Observe that *U* ^′^ consists of exactly (elements corresponding to) all of the red edges and that the solutions for the Set Cover instance (*U*, {*A*_*i*_}_*i*_) are in one-to-one correspondence to solutions for instance 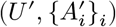.

Next, we map each set 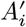 to a walk *W*_*i*_ in the graph consisting only of red edges in a way that we can connect the walks *W*_*i*_ into one walk in *G* using only blue edges. For every set 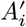 we list all the light-red edges which were mapped to an element in this set. We construct a walk *W*_*i*_ in the following manner. We start with the edge corresponding to the first element in *A*_*i*_ and iteratively append edges corresponding to other elements of *A*_*i*_ to *W*_*i*_, connected by a path of dark-red edges in *H*_*R*_. Namely, suppose that *W*_*i*_ ends with the *j*-th element of *A*_*i*_ and we want to add the (*j* + 1)-th. The walk ends at some vertex *v*_*j*_ and the next edge starts at possibly different vertex *u*_*j*+1_. We take the path *P*_*j*_ from *v*_*j*_ to *u*^*j*+1^ in the dark-red Hamiltonian cycle *H*_*R*_. Then we append path *P*_*j*_ and the (*j* + 1)-th edge (corresponding to the (*j* + 1)-th element of *A*_*i*_) to *W*_*i*_, thus extending this walk by one element from the set. At the end, we append the whole dark-red Hamiltonian cycle *H*_*R*_ to *W*_*i*_. This way we obtain a walk which contains precisely the red edges corresponding to elements in *A*^′^. Note also that *W*_*i*_ is polynomially large with respect to the size of the Set Cover instance as it contains at most (*n* + 1)|*A*_*i*_| + *n* elements.

It remains to show that we can connect these walks with walks of blue edges. We can use the same trick as with connecting edges within the set. We take the subpath of the blue Hamiltonian cycle *H*_*B*_ from the end of the *i*-th walk to the beginning of the *i* + 1-th and concatenate the two walks together. Eventually, we get a walk *W* such that every maximal red subwalk of *W* is *W*_*i*_ for some *i*. Therefore, the reduction preserves Set Cover solutions exactly, with the same objective value (i.e., every solution for the Set Cover instance corresponds to a solution for MaskMinNumRuns on the instance from the reduction, and vice versa, and moreover, the number of selected subsets *A*_*i*_ equals the number of selected red subwalks, or the number of runs of ones). This concludes the polynomial-time reduction from Set Cover to the graph formulation of MaskMinNumRuns, which implies that the NP-hardness of approximation by Theorem G.2.

The main downside of the proof is that the superstring resulting from the reduction would hardly be computed by any reasonable superstring algorithm, even on the same set of *k*-mers as in the reduction (moreover, a set of *k*-mers with two edge-disjoint Hamiltonian cycles in its de Bruijn graph may not occur in practice). Still, the hardness proof justifies the usage of ILP solvers for MaskMinNumRuns outlined in Appendix H. We leave it as an open question whether or not for a *k*-mer superstring computed by a particular algorithm, e.g., local or global greedy, it is possible to solve MaskMinNumRuns in polynomial time.

## H Minimizing the number of runs using integer linear programming

Despite the NP-hardness result in Appendix G, we provide a method that, for a given superstring *S* and a set of *k*-mers, finds a mask for *S* that has the minimum number of runs of ones. As we describe next, this problem can be solved by integer linear programming (ILP) for which there are practically efficient solvers. To make it even more efficient, we first run a simple greedy heuristic, which reduces the ILP size significantly.

### Greedy heuristic

We start by observing that any position 0 ≤ *i* ≤ |*S*| − *k* in superstring *S* is of one of the following two types:

**Type-0:** A ghost *k*-mer starts at *i*, i.e., string *S*[*i* ∶ *i* + *k*] and its reverse complement (RC) are not in the set of *k*-mers. Any mask for *S* must have a 0 at position *i*. (The last *k* − 1 positions in *S* are of type 0.)

**Type-?:** A *k*-mer (or the RC of a *k*-mer) starts at position *i*. A mask for *S* may have 1 or 0 at position *i*.

Type-0 positions naturally split the set of indexes {0, …, |*S*| −*k*} into maximal intervals of type-? positions that we just call *intervals* for brevity.

The optimal mask *M* that we will construct starts with either 0 or ? at any position *i*, depending on the type of *i*. We use two simple observations: Any *k*-mer needs to be represented just once (possibly as its RC), and once an optimal mask has at least one position set to 1 inside an interval, the whole interval may consist of 1s only. Thus, for each interval, we just need to decide whether to set 1 or 0 in the whole interval. The greedy heuristic then goes as follows:

1. For each interval, if there is a *k*-mer a such that a (or its RC) does not appear in any other interval, this interval must contain a 1 in any mask and therefore, we set 1 in the constructed mask for all positions in the whole interval._1_
2. After that, if there is an interval *I* not already set to 1 such that every *k*-mer inside *I* is already represented by an interval set to 1 by the previous case, we set interval *I* to 0 (i.e., all positions in *I* are set to 0 in the mask).

We are left with a set of *undecided intervals* (those not set to 0 or 1) and *non-represented k-mers* (there is no position already set to 1 that represents such a *k*-mer). Note that an undecided interval must contain a non-represented *k*-mer and a non-represented *k*-mer appears in at least two undecided intervals. Furthermore, a non-represented *k*-mer does not appear in an interval falling in one of the two cases of the greedy heuristic. Our experiments on the streptococcus genomes and pangenomes show that, undecided intervals are still present after we run the greedy heuristic, but their number is significantly smaller than the total number of intervals.

### Integer linear program

To complete the computation of the optimal mask, we construct an ILP model as follows: For each undecided interval *I*, there is a binary variable *x*_*I*_, and for each non-represented *k*-mer *a*, we add a constraint that the sum of *x*_*I*_*’*s over all undecided intervals containing a is at least one. The objective is to minimize the sum of *x*_*I*_*’*s over all undecided intervals *I*. An optimal solution immediately gives an optimal mask by setting (all positions in) each interval *I* to *x*_*I*_. We thus obtain the following ILP model, where *𝒰* is the set of undecided intervals:

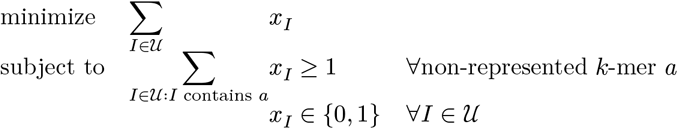

We remark that this ILP is similar to the one for Set Cover; indeed, minimizing the number of runs is similar in essence to solving Set Cover as also shown by the reduction in Appendix G.

## Footnotes

1 In the actual implementation, we take a simpler approach: Count the number of occurrences of each canonical *k*-mer, and only set 1 in an interval if there is a *k*-mer that appears just once in the whole superstring. In this way, possibly less intervals fall into the first case, namely, intervals *I* containing a *k*-mer a more than once such that a does not appear in another interval (and there is no *k*-mer in *I* that appears just once).

## Notes

### Competing Interest Statement

The authors have declared no competing interest.

